## Supplementary figures and images for "Resolving the origins of secretory products and anthelmintic responses in a human parasitic nematode at single-cell resolution"

### Fig1-S1

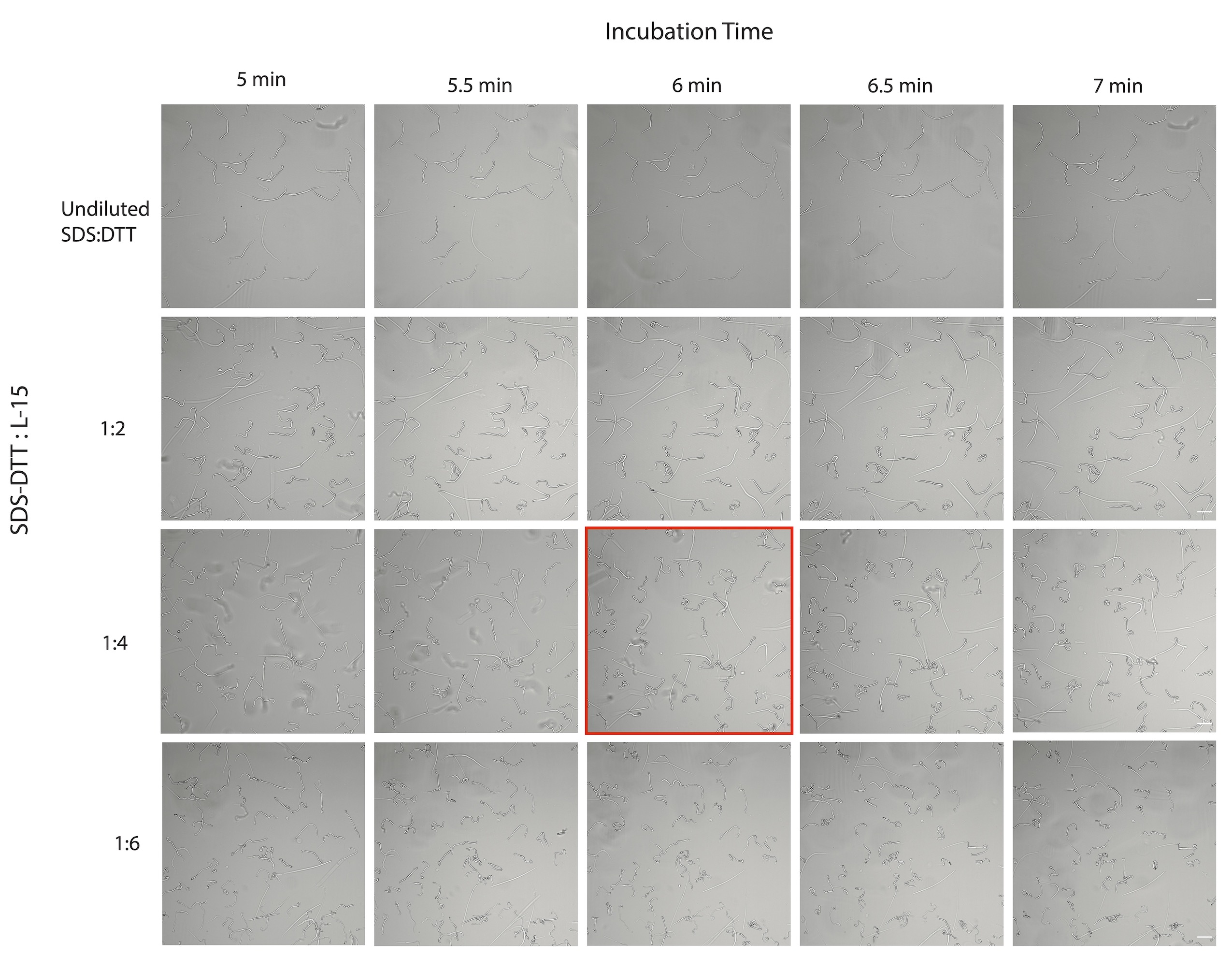

### Fig2-S1

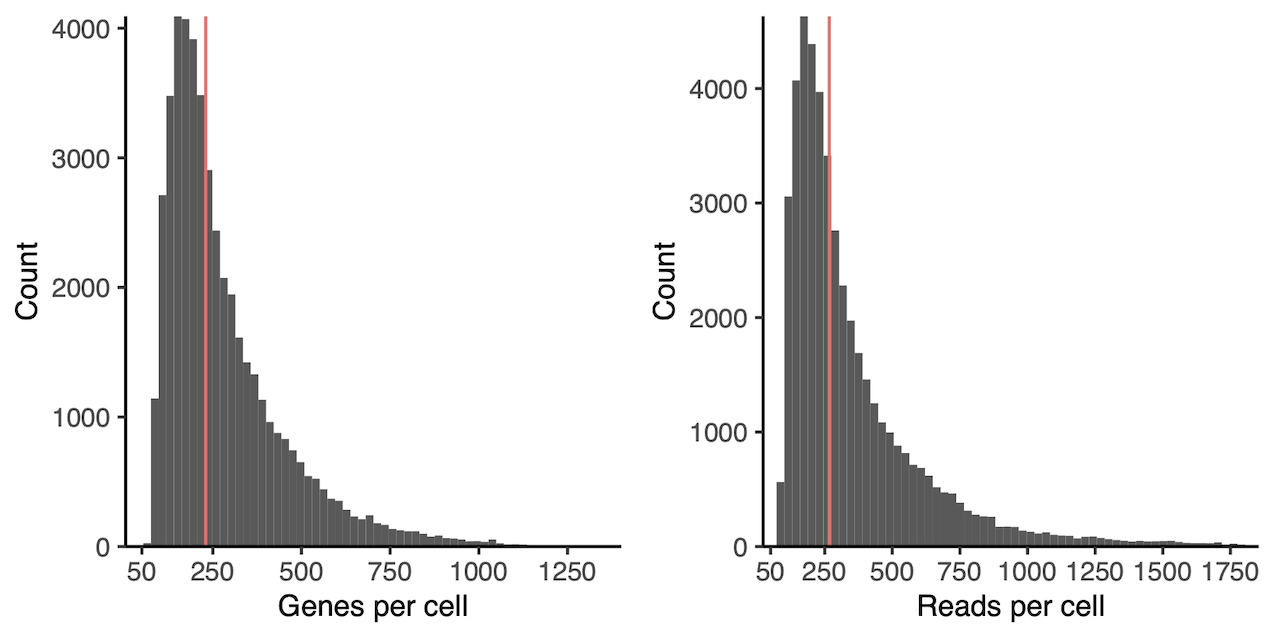

### Fig2-S2

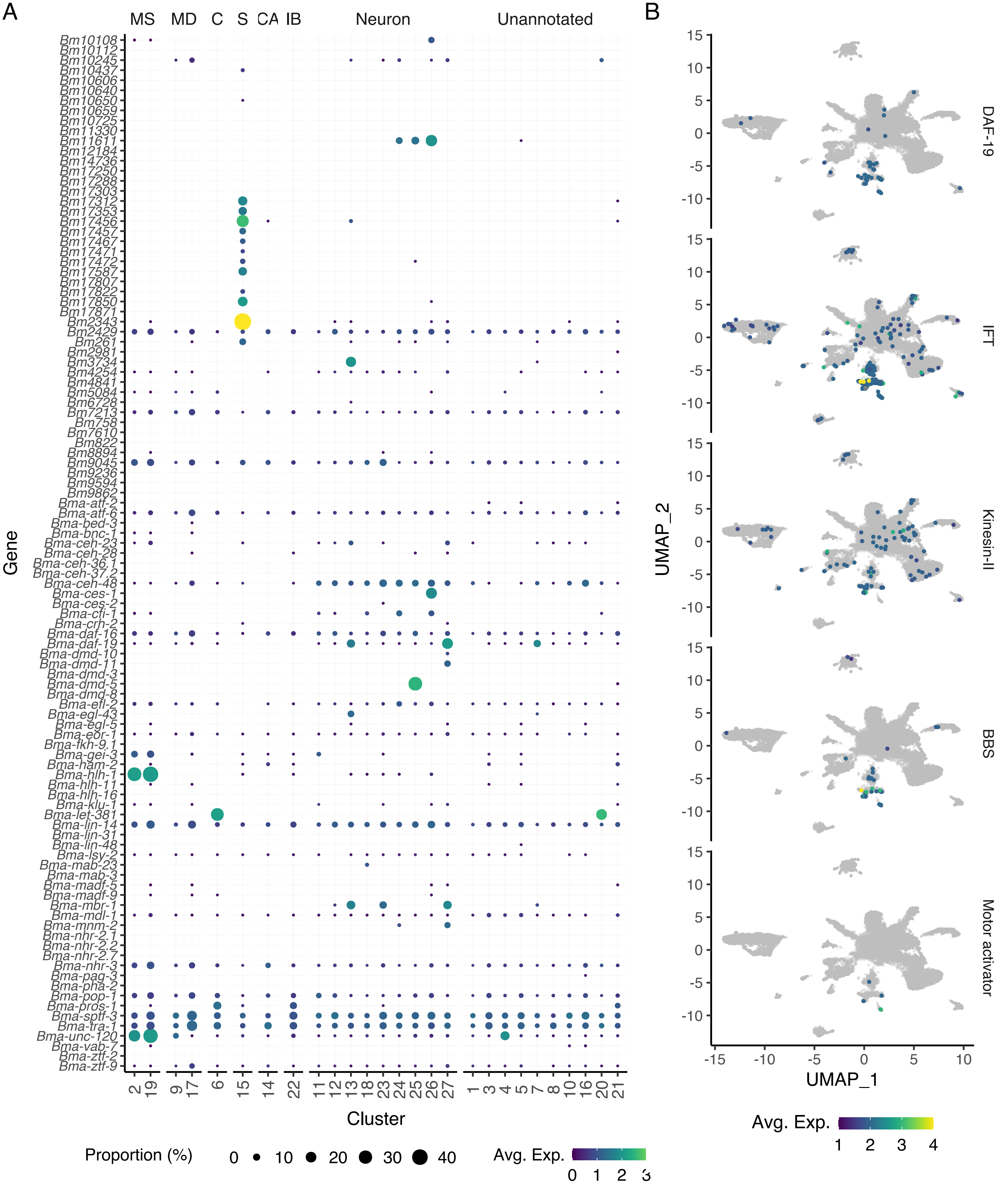

### Fig2-S3

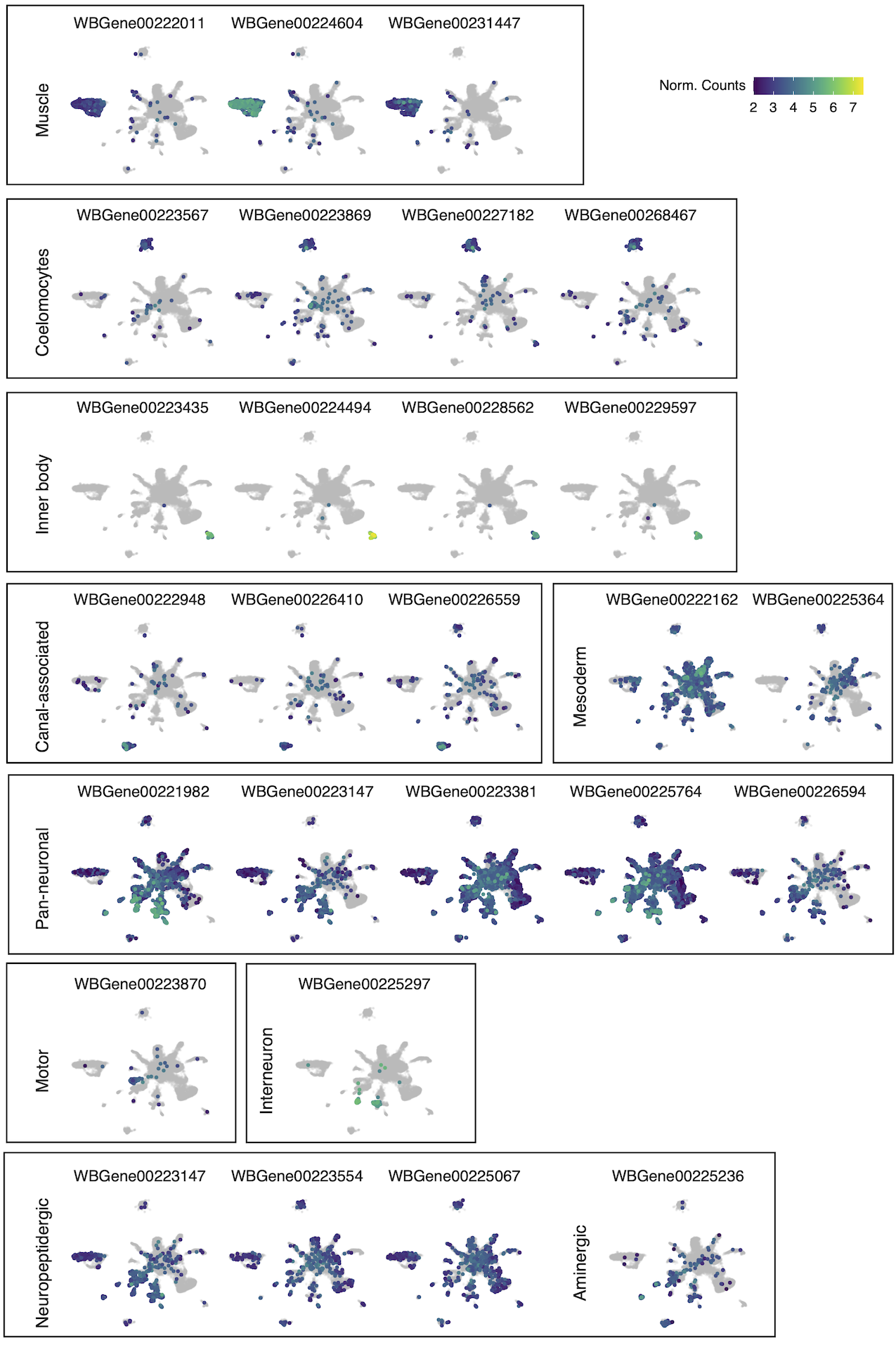

### Fig2-S4

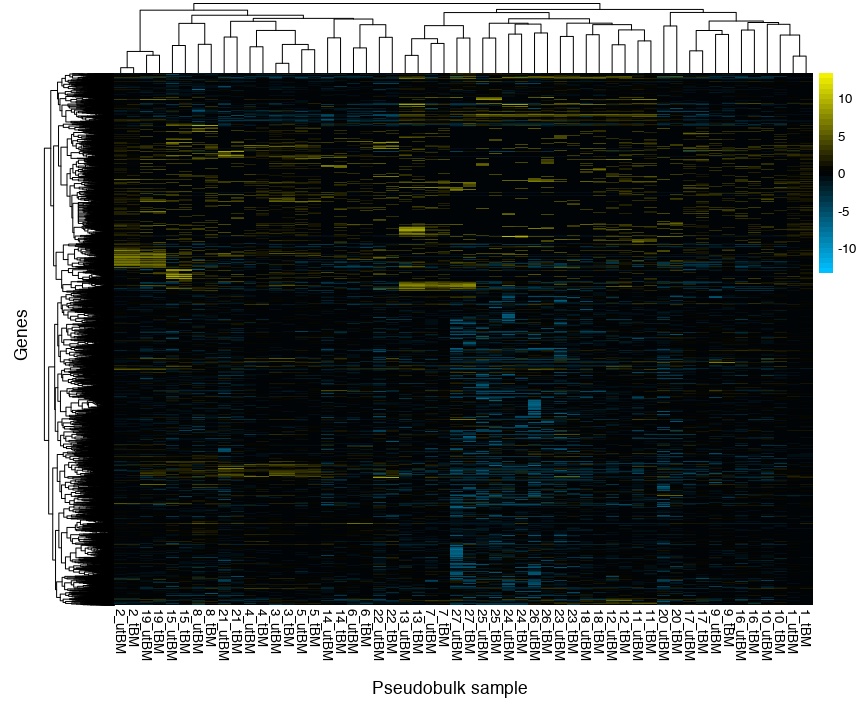

### Fig2-S5

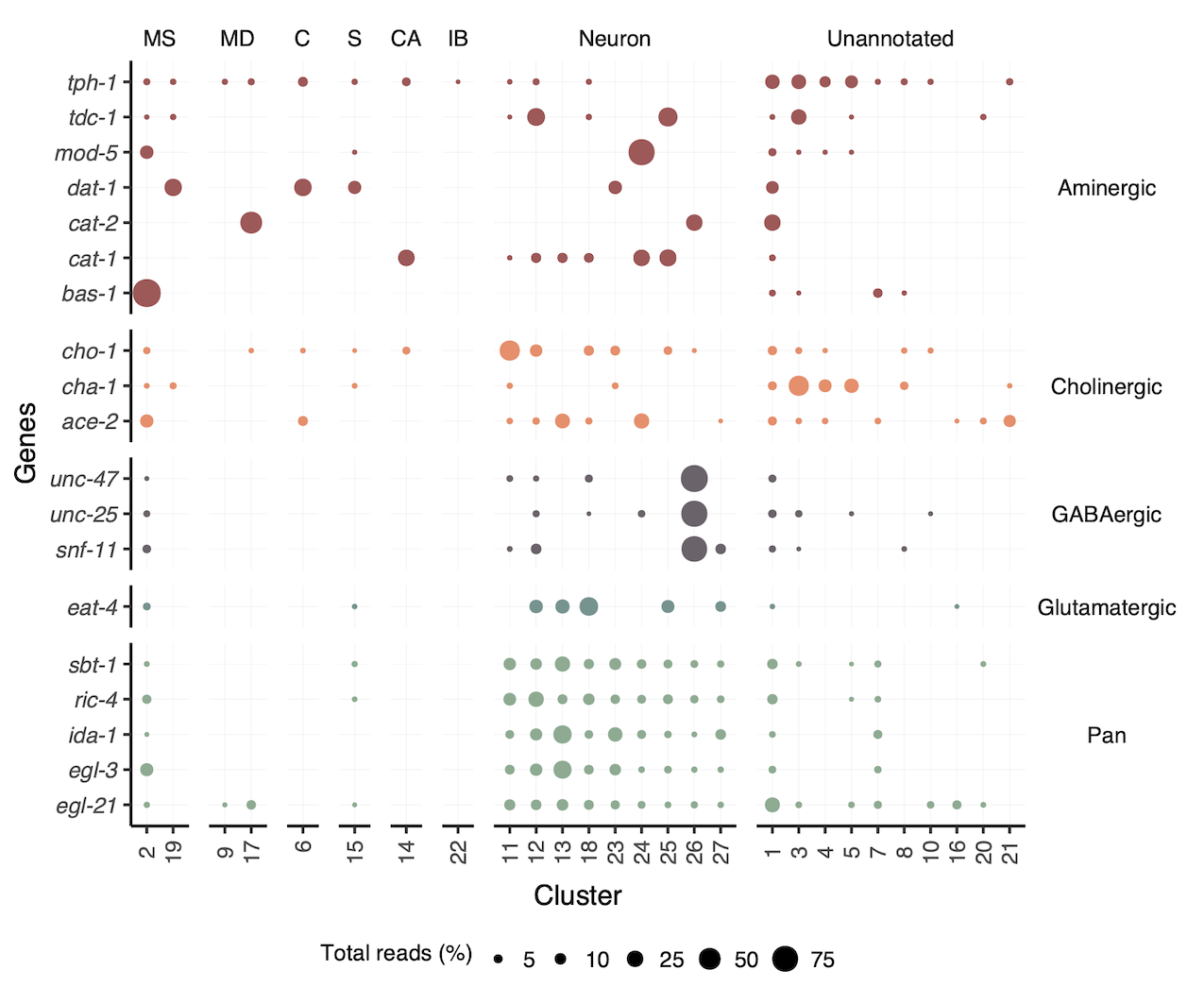

### Fig2-S6

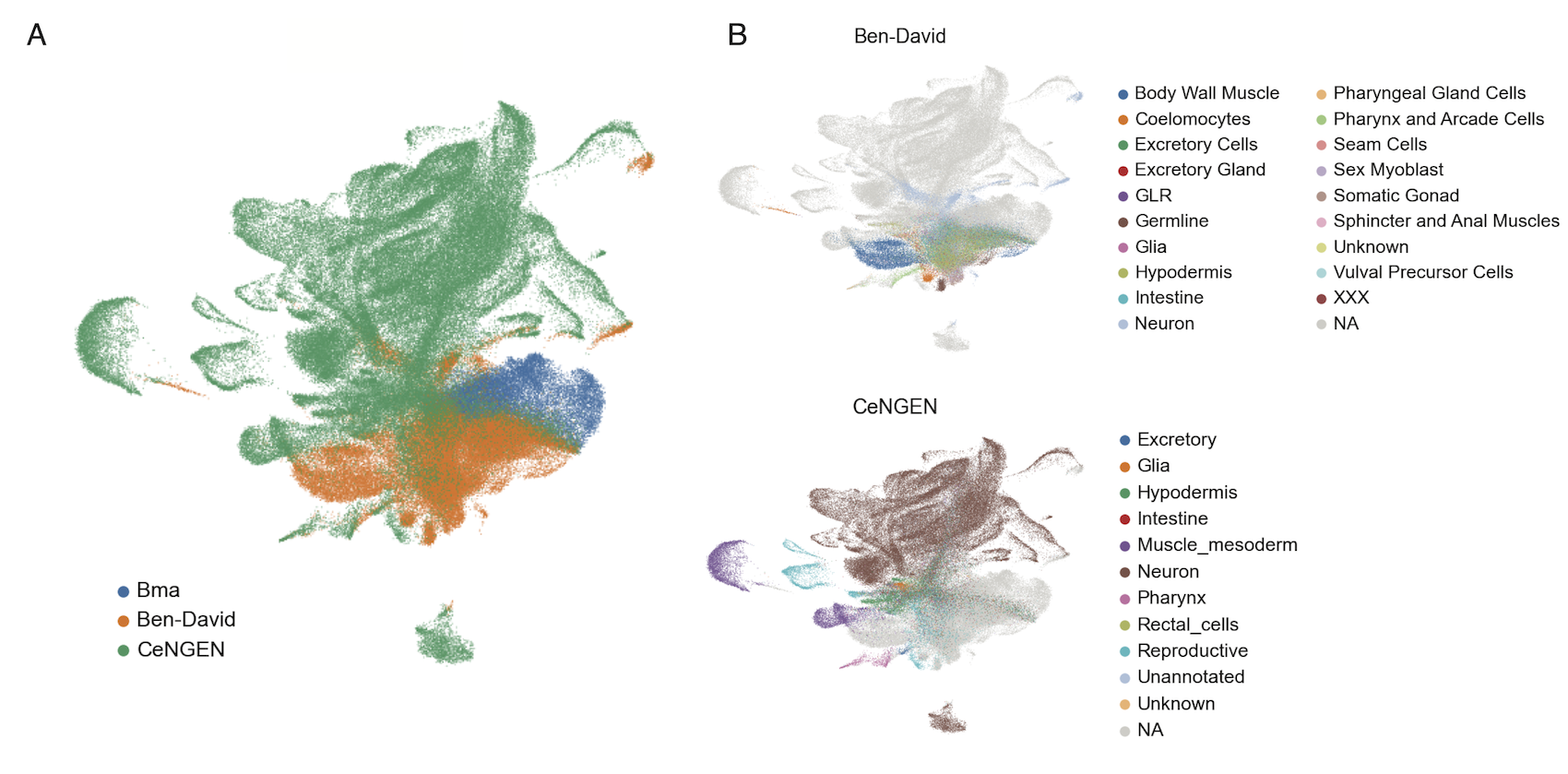

### Fig3-S1

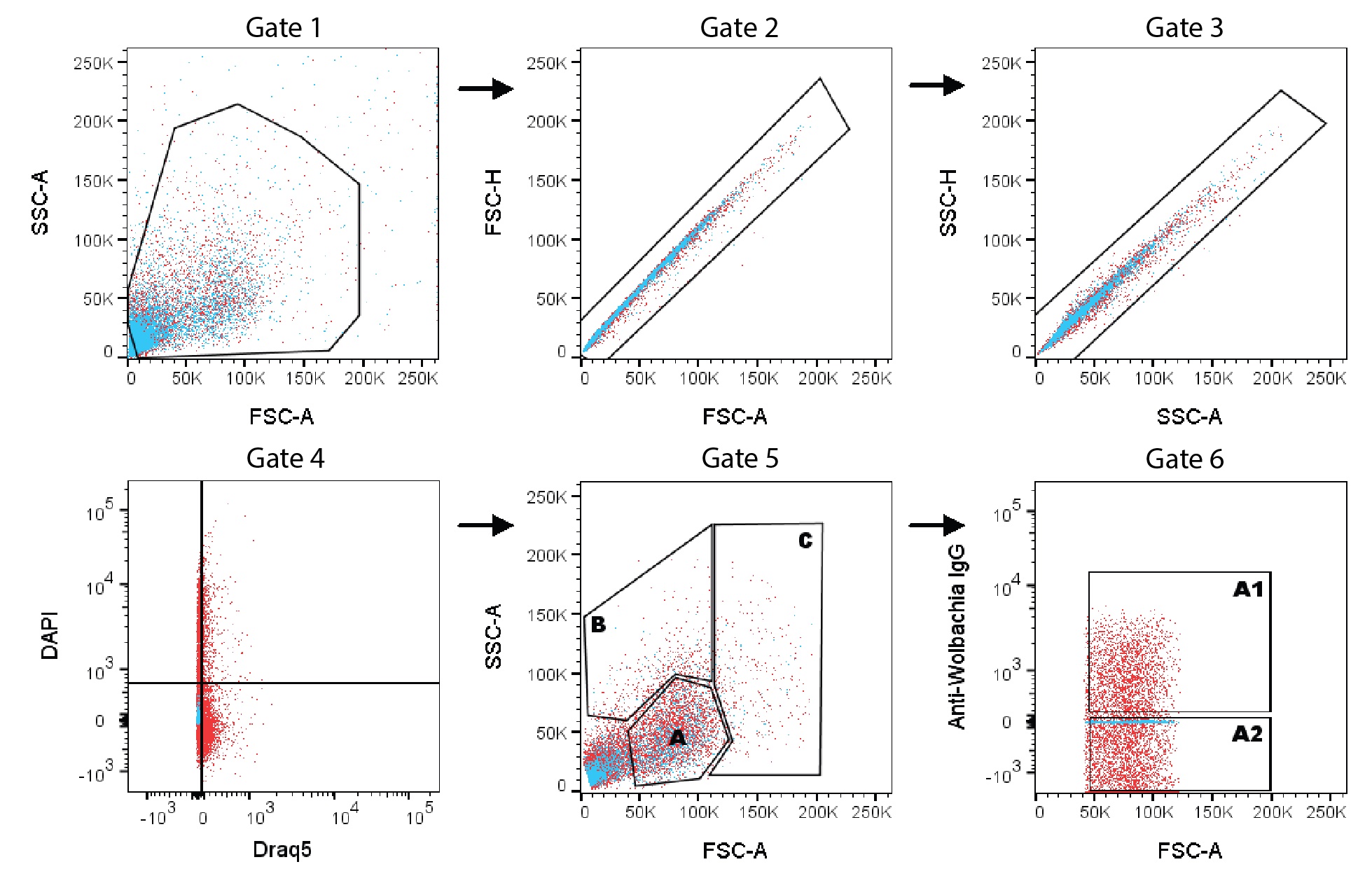

### Fig3-S2

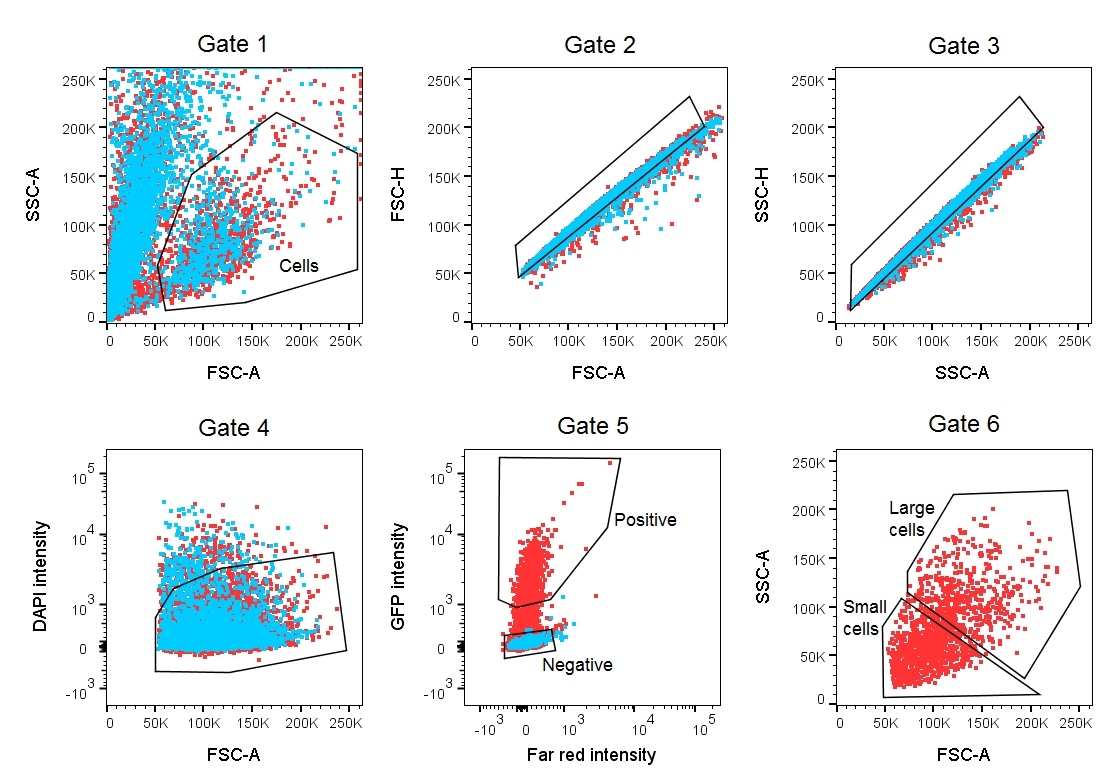

### Fig3-S3

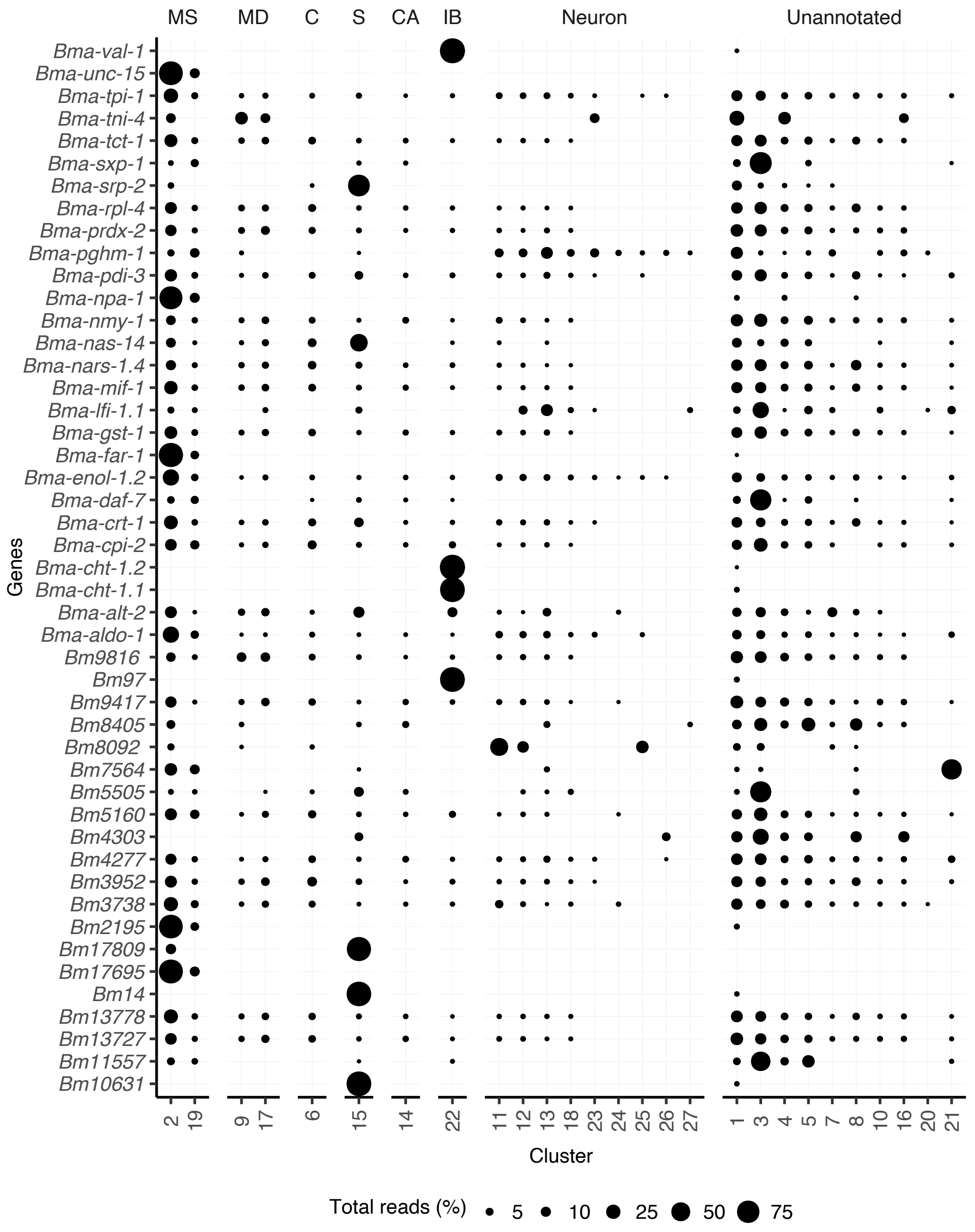

### Fig4-S1

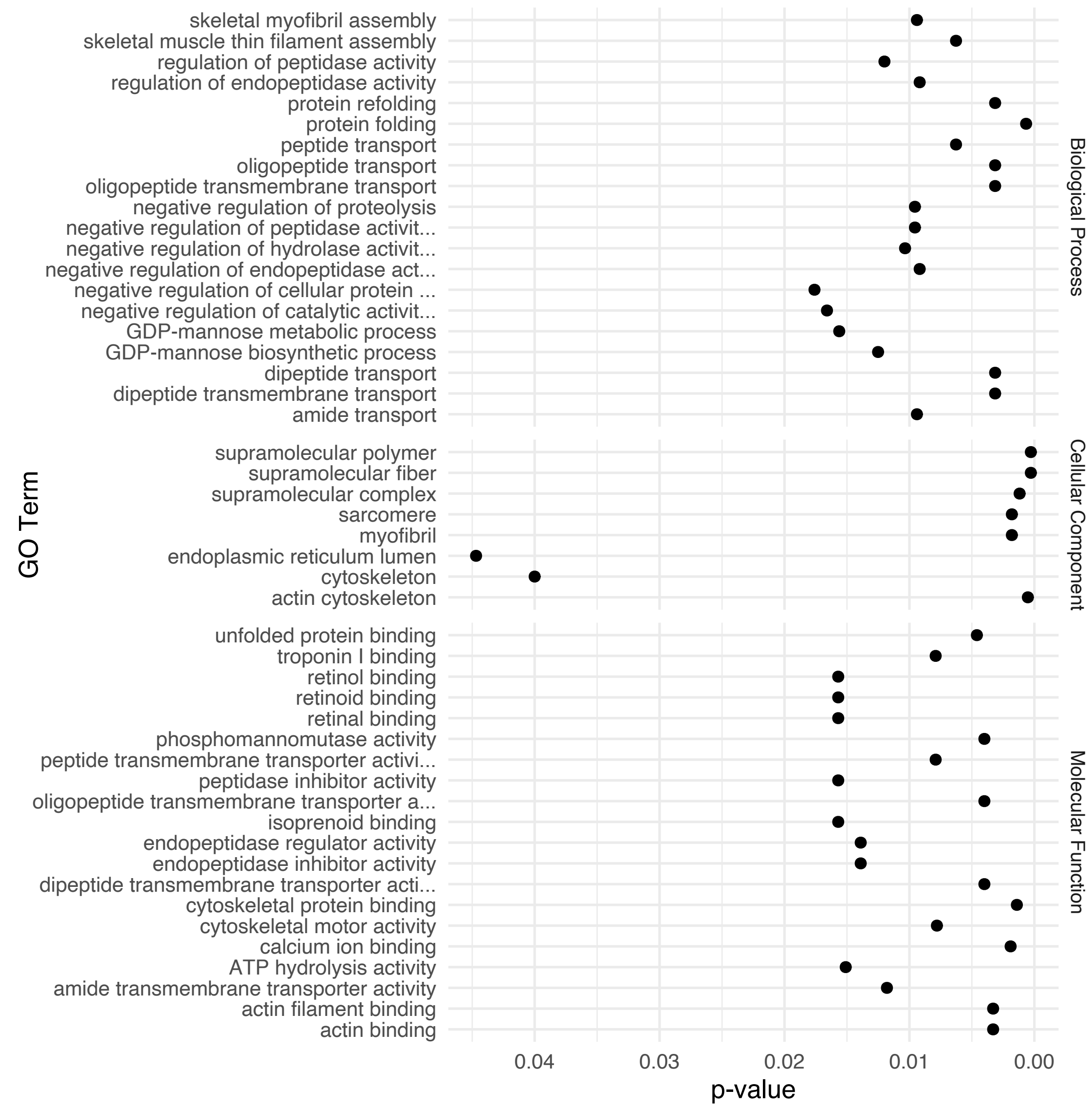

### Fig5-S1

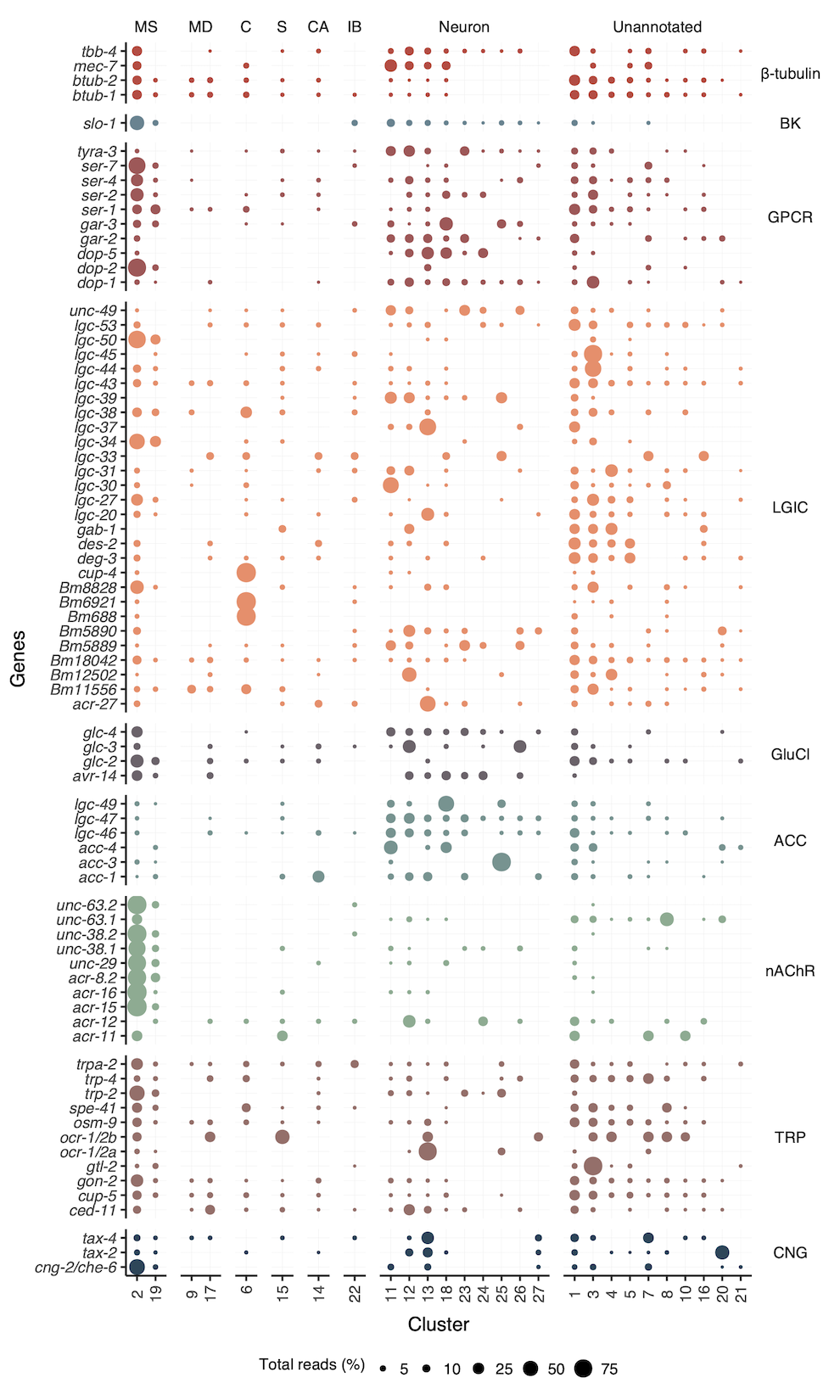

### Fig7-S1

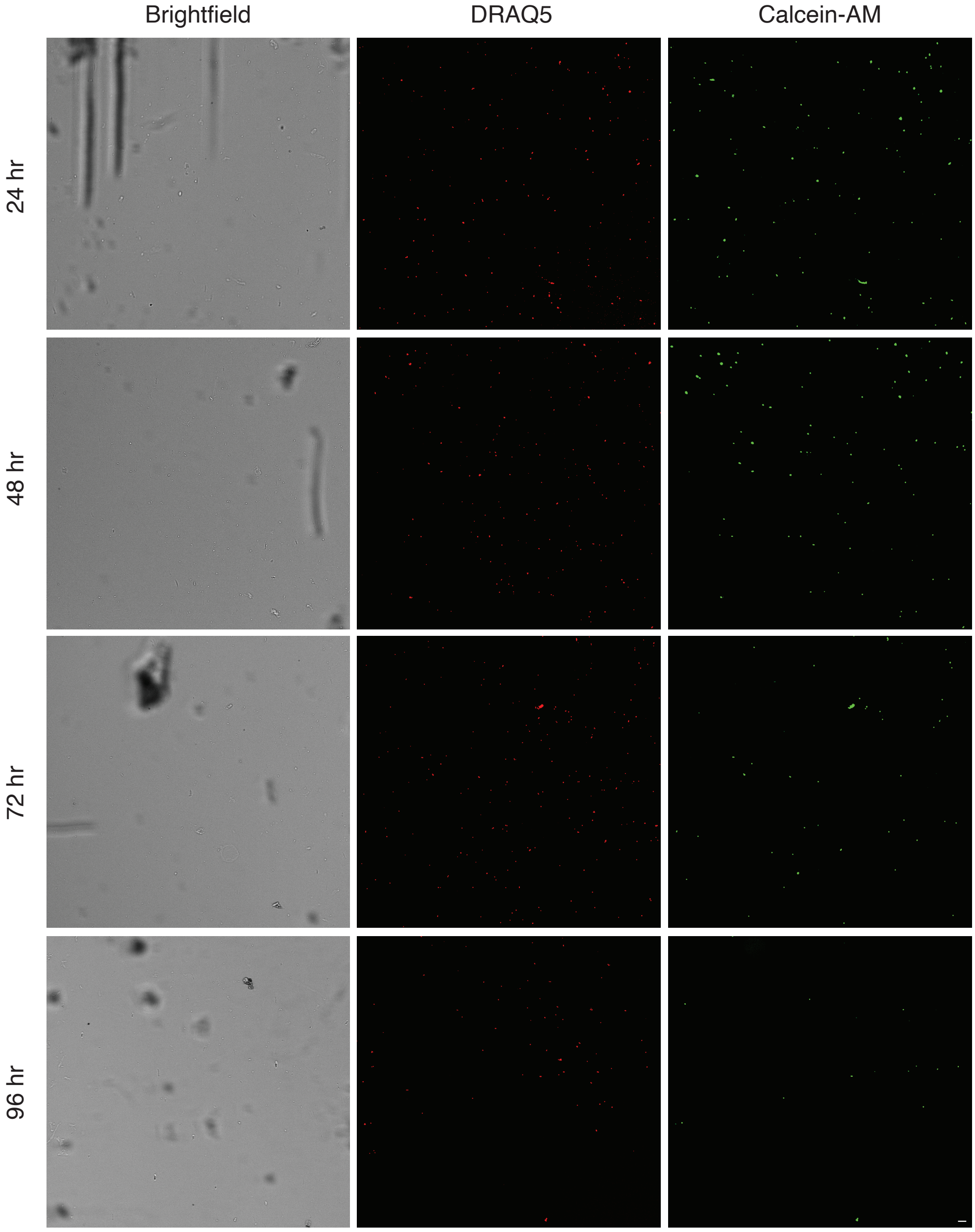

### Fig7-S2

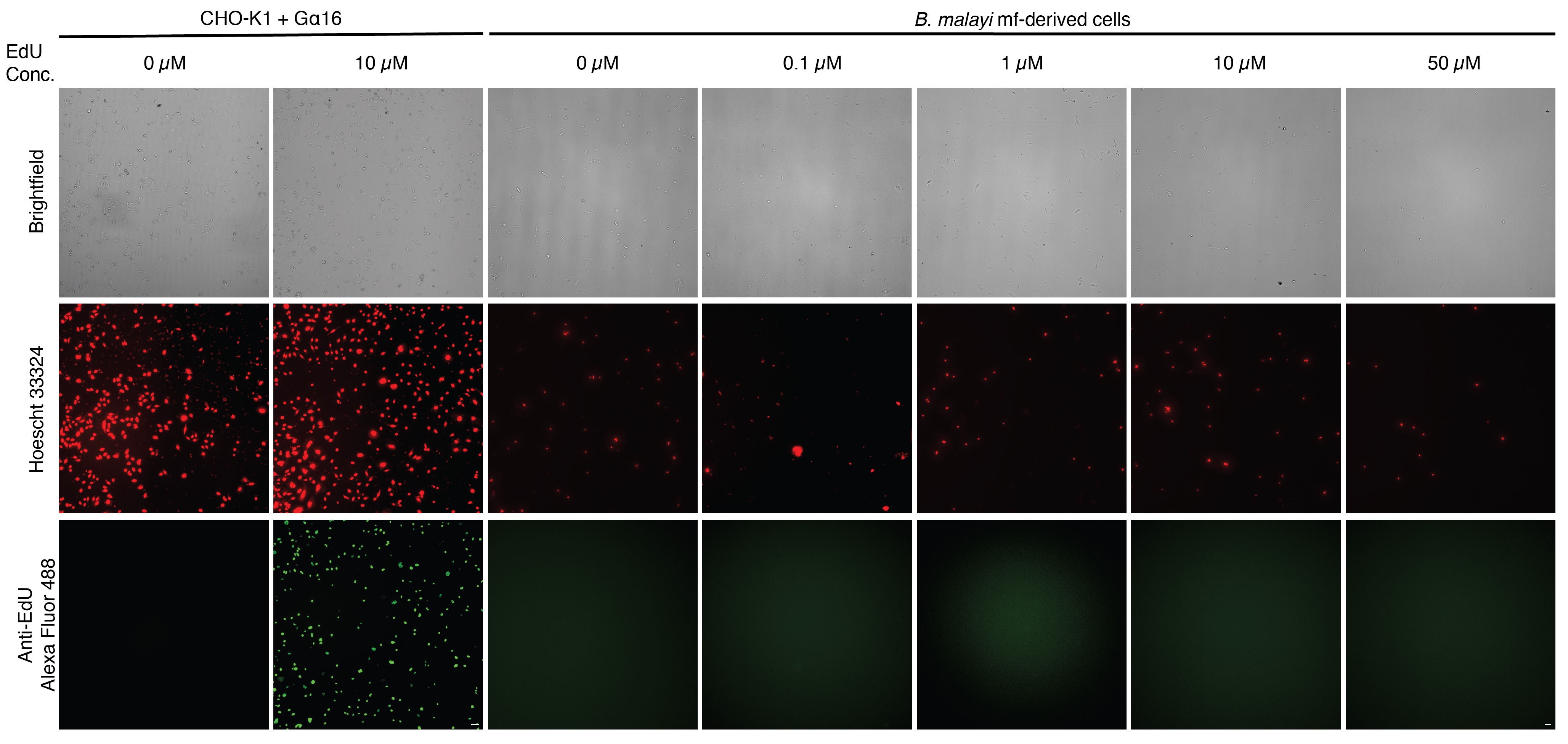
