## Supplementary material for "Resolving the origins of secretory products and anthelmintic responses in a human parasitic nematode at single-cell resolution": Fig1-D1

date	species	stage	measurement	staining	input	total_cells	viable_cells	units	notes

20201110	Bpa	mf	FACS	draq5/dapi	300000	7505	1644

20201110	Bpa	mf	FACS	draq5/dapi	120000000	17811	4536

20201110	Bpa	mf	CountessII	draq5/dapi	300000	167000	8785	cells/mL	pre-sort

20201110	Bpa	mf	CountessII	draq5/dapi	120000000	1610000	29350	cells/mL	pre-sort

20201110	Bpa	mf	CountessII	draq5/dapi	300000	44000	0	cells/mL	post-sort

20201110	Bpa	mf	CountessII	draq5/dapi	120000000	88000	0	cells/mL	post-sort

20201117	Bma	mf	FACS	draq5/dapi	120000000	4040000	500000	cells/mL

20201117	Bma	mf	CountessII	draq5/dapi	120000000	49800	49830	cells/mL	post-sort

20201117	Bma	mf	hemocytometer	NA	120000000	1870000	NA	cells/mL

20201215	Bma	mf	CountessII	NA	120000000	633000	NA	cells/mL

20201215	Bma	mf	hemocytometer	NA	120000000	2070000	NA	cells/mL

20201215	Bma	mf	FACS	draq5/dapi	120000000	669812	71292	cells/mL
